# Impact of Solvent Extraction on Bioactive Properties of *Lavandula* × *intermedia*: Antibacterial, Antioxidant and Phenolic Content Analysis

**DOI:** 10.64898/2026.07.11.737895

**Authors:** Gülcan Gürses

## Abstract

*Lavandula × intermedia* (lavandin) is an economically important hybrid valued for its high essential oil yield; however, studies linking solvent polarity to its bioactivity and phenolic composition remain limited. This study evaluated the effects of nine extraction solvents on the antibacterial and antioxidant activities of *L. × intermedia*, determined total phenolic content (TPC), and examined the relationship between chemical composition and biological activity. Aerial parts were extracted by maceration. Antibacterial activity was assessed using the broth microdilution method against four bacterial strains, while antioxidant capacity was measured by the DPPH assay. TPC was determined using the Folin–Ciocalteu method, and phenolic compounds were analyzed via LC–MS/MS. Results showed that solvent polarity significantly influenced bioactivity. Diethyl ether extracts exhibited the highest TPC (267.65 mg GAE/g), strongest antioxidant activity (IC_50_: 72.26 µg/mL), and notable antibacterial effects (MIC: 125 µg/mL), especially against Gram-positive bacteria. A strong positive correlation (*r* = 0.887) was observed between phenolic content and antimicrobial activity. LC–MS/MS identified key compounds, including fumaric acid, resveratrol, and hydroxycinnamic acid. Overall, moderate-polarity solvents such as diethyl ether and methanol were most effective, highlighting *L. × intermedia* as a promising natural source for pharmaceutical and nutraceutical applications.

## Introduction

The genus *Lavandula* (Lamiaceae), comprising 39 species and over 400 cultivars, holds significant economic value in medicine and perfumery (Ez Zoubi et al. 2020). Native to the Mediterranean, Africa, and Asia, these aromatic plants are increasingly recognized for their broad-spectrum therapeutic properties, including antioxidant, antimicrobial, and neuroprotective effects (Koulivand et al. 2013; Batiha et al. 2023; Pokajewicz et al. 2023a; Cardia et al. 2021). These attributes are primarily driven by the complex phytochemical matrix within their essential oils and extracts (Cardia et al. 2021).

Commonly utilized species like *L. angustifolia, L. latifolia, L. stoechas*, and the hybrid *L. × intermedia* are cultivated globally for applications ranging from aromatherapy and cosmetics to the food and detergent industries (Basch et al. 2004; Zuzarte et al. 2009). Their long history in folk medicine is supported by anti-inflammatory, analgesic, sedative, and antimicrobial properties (Giovannini et al. 2016; Hassiotis 2010; Gilani et al. 2000), attributed to a rich content of phenolic acids, flavonoids, and volatile terpenoids (Moon et al. 2006; Jianu et al. 2013; Lahkimi et al. 2020; Ciocarlan et al. 2021).

*Lavandula × intermedia* Emeric ex Loisel. (lavandin), a sterile hybrid of *L. angustifolia* and *L. latifolia*, is valued for its robustness and high essential oil yield, which can quintuple that of *L. angustifolia* (Pokajewicz et al. 2022; Lis-Balchin 2002). While its bioactivity is documented, potency varies by geography and extraction method (Truong and Mudgil 2023; Roller et al. 2009; Mesic et al. 2021; Valková et al. 2021). Few studies have comprehensively evaluated how solvent polarity dictates phenolic composition and bioactivity using advanced methods like LC–MS/MS, a critical gap for optimizing the plant of potential (Park et al. 2019; Danh et al. 2013).

Traditional literature emphasizes terpenes like linalool and camphor in essential oils (Woronuk et al. 2011; Prusinowska and Smigielski 2014). However, the non-volatile polyphenols found in polar extracts (methanol, ethanol, water) significantly contribute to anti-inflammatory and radical scavenging activities (Batiha et al. 2023). Since existing research often focuses on limited solvent ranges, a systematic comparison across a broad polarity spectrum for *L. × intermedia* is missing, hindering optimized commercial protocols.

To address this, we evaluated nine solvents (apolar to polar) to correlate antibacterial and antioxidant activities with specific LC-MS/MS fingerprints. Our hypothesis is that moderate-polarity solvents, such as diethyl ether and methanol, maximize phenolic diversity and bioactivity. We quantified Total Phenolic Content (TPC), characterized chemical profiles via LC–MS/MS, and statistically analyzed the relationships between solvent polarity and biological efficacy to establish an optimized extraction protocol.

## Materials and Methods

### Preparation of plant extracts

*Lavandula × intermedia* samples were collected from Aligor–Suruç (Şanlıurfa, Türkiye) in July 2025. Locality: 37°01’42.1”N, 38°26’22.5”E (WGS-84 datum; ± 10 m), Şanlıurfa Province, Türkiye. The plant material was taxonomically identified by Dr. Mehmet Maruf Balos, Department of Biology, Harran University. A voucher specimen (Collector No: M.Balos 5722) was deposited in the Harran University Faculty of Pharmacy Herbarium (HUEH). Harvested plants were air-dried at ambient temperature for seven days and pulverized. Extraction was performed via maceration in nine solvents (ethanol, water, diethyl ether, methanol, hexane, chloroform, dichloromethane, acetone, and ethyl acetate) at a 1:10 (w/v) ratio. Briefly, 10 g of powder was immersed in 100 mL of solvent for 72 hours at 25 ± 2°C with daily agitation. After maceration, extracts were filtered through 0.45 µm membranes, diluted with distilled water to 16,000 µg/mL, and sterilized via 0.2 µm Acrodisc syringe filters. Samples were stored at –20°C until further use.

### Bacterial strains and inoculum preparation

Gram-positive (*Staphylococcus aureus* ATCC 29213, *Enterococcus faecalis* ATCC 29212) and Gram-negative (*Escherichia coli* ATCC 25922, *Pseudomonas aeruginosa* ATCC 27853) strains were used. Isolates were cultured on blood agar and eosin methylene blue (EMB) agar at 37°C for 18–24 hours under aerobic conditions. Strains were subcultured for experimental use and stored in Tryptic Soy Broth (TSB) with glycerol at −20°C. For assays, colonies from fresh cultures were suspended in Mueller–Hinton Broth (MHB) and adjusted to a 0.5 McFarland standard (approximately 1.5 × 10^8^ CFU/mL) using a densitometer.

### Determination of minimum inhibitory concentration (MIC)

The minimum inhibitory concentrations (MICs) of the plant extracts were determined using the broth microdilution method in 96-well microplates, following the guidelines of the Clinical and Laboratory Standards Institute (CLSI) with minor modifications. Stock solutions of each extract were prepared in sterile distilled water at a final concentration of 16 mg/mL. In each well of a sterile 96-well microplate, 100 µL of Mueller–Hinton Broth (MHB) was dispensed. Then, 100 µL of the plant extract was added to the wells of the first column, followed by two-fold serial dilutions across the plate to obtain final concentrations of 8000-62.5 µg/mL. The bacterial suspensions were first adjusted to the 0.5 McFarland (∼ 1.5 × 10^8^ CFU/mL) and then diluted 1:15 in fresh MHB to prepare a working suspension. Finally, 10 µL of this working suspension was inoculated into each well. This procedure ensured that the final bacterial concentration in the wells approximated the standard value of 5 × 10^5^ CFU/mL. Gentamicin was selected as a broad-spectrum positive control, adjusted to 0.06–16 μg/mL. Microplates were incubated at 37°C for 24 hours under aerobic conditions. MIC was defined as the lowest concentration of extract at which no visible bacterial growth (turbidity) was observed compared to the controls. All tests were performed in triplicate.

### Determination of antioxidant activity (DPPH assay)

The antioxidant capacity was evaluated via the DPPH radical scavenging assay following Blois (1985) and Necip and Durgun (2022) with minor modifications. Extract stock solutions (1.00 mg/mL) were prepared, and aliquots (10, 20, 30 µL) were diluted to 1.0 mL with distilled water, then adjusted to 2.0 mL with absolute ethanol. Subsequently, 0.5 mL of fresh DPPH solution (in ethanol) was added. Mixtures were vortexed and incubated in the dark at room temperature for 30 minutes. Absorbance was measured at 517 nm using a UV–Vis spectrophotometer. Ethanol served as the negative control. The scavenging activity percentage was calculated as follows:

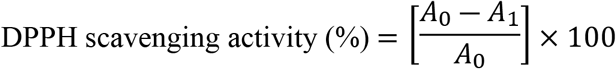

where *A*_0_ is the absorbance of the control solution and *A*_1_ is the absorbance of the test solution containing the extract.

### Determination of total phenolic content

Total phenolic content was determined using a modified Folin–Ciocalteu method (Singleton et al. 1999). Briefly, 50 µL of extract was mixed with 1150 µL of distilled water and 25 µL of Folin–Ciocalteu reagent. After a 3-minute incubation, 75 µL of 7.5% (w/v) Na_2_CO_3_ was added. The mixture was vortexed and incubated in the dark at room temperature for 2 hours. Absorbance was measured at 760 nm using a UV–Vis spectrophotometer. TPC was calculated via a gallic acid calibration curve and expressed as milligrams of gallic acid equivalents per gram of extract (mg GAE/g extract) (Amangeldinova et al. 2025).

### Determination of phenolic compounds by LC–MS/MS

The phenolic profile of *L. × intermedia* methanol extracts was analyzed using a Shimadzu LCMS-8030 triple quadrupole mass spectrometer coupled with a Shimadzu LC-20ADXR UHPLC system. Separation was performed on a SUPELCO Discovery C18 column (50 × 2.1 mm, 5 µm) at 40°C. The mobile phase consisted of 0.1% formic acid in water/methanol mixtures (Solvent A: 20/80; Solvent B: 80/20 v/v) with a 15-minute gradient program at a flow rate of 0.4 mL/min. Detection was carried out in Multiple Reaction Monitoring (MRM) mode with ESI (positive/negative). The method was validated per ICH Q2(R2) guidelines. All analytes showed excellent linearity (*R*^2^: 0.9980–0.9999) within the 10–2500 µg/L range. LOD and LOQ values ranged from 2.76–38.5 µg/L and 9.20–128.2 µg/L, respectively. Quantification was performed using independent external calibration curves.

## Results and Discussion

### Antibacterial activity of *Lavandula × intermedia* extracts

The antibacterial efficacy of *L. × intermedia* extracts varied by solvent type (Table 1). Diethyl ether exhibited the most potent activity, with the lowest MIC values (125 µg/mL) against *S. aureus, E. coli*, and *E. faecalis*. Similarly, methanol extracts showed strong inhibition, with MIC values of 125 µg/mL for *S. aureus* and *E. faecalis*, and 250 µg/mL for other strains (Fig. 1). In contrast, the water extract demonstrated the weakest activity, particularly against *E. coli* (1000 µg/mL), highlighting the limited efficacy of polar aqueous extraction. Generally, *P. aeruginosa* was the most resistant strain. These findings confirm that solvents of moderate polarity are the most efficient for isolating the synergistic combination of phenolic compounds responsible for antimicrobial action. Gram-positive bacteria possess a thick, single-layered peptidoglycan cell wall, facilitating the passage of bioactive molecules, whereas Gram-negative bacteria are protected by a complex outer membrane and a lipopolysaccharide (LPS) layer (Nikaido 2003). These results align with previous studies reporting lavender’s broad antibacterial activity against Gram-positive strains and MRSA (Truong and Mudgil 2023; Roller et al. 2009; Sasaki et al. 2015).

**Table 1.**
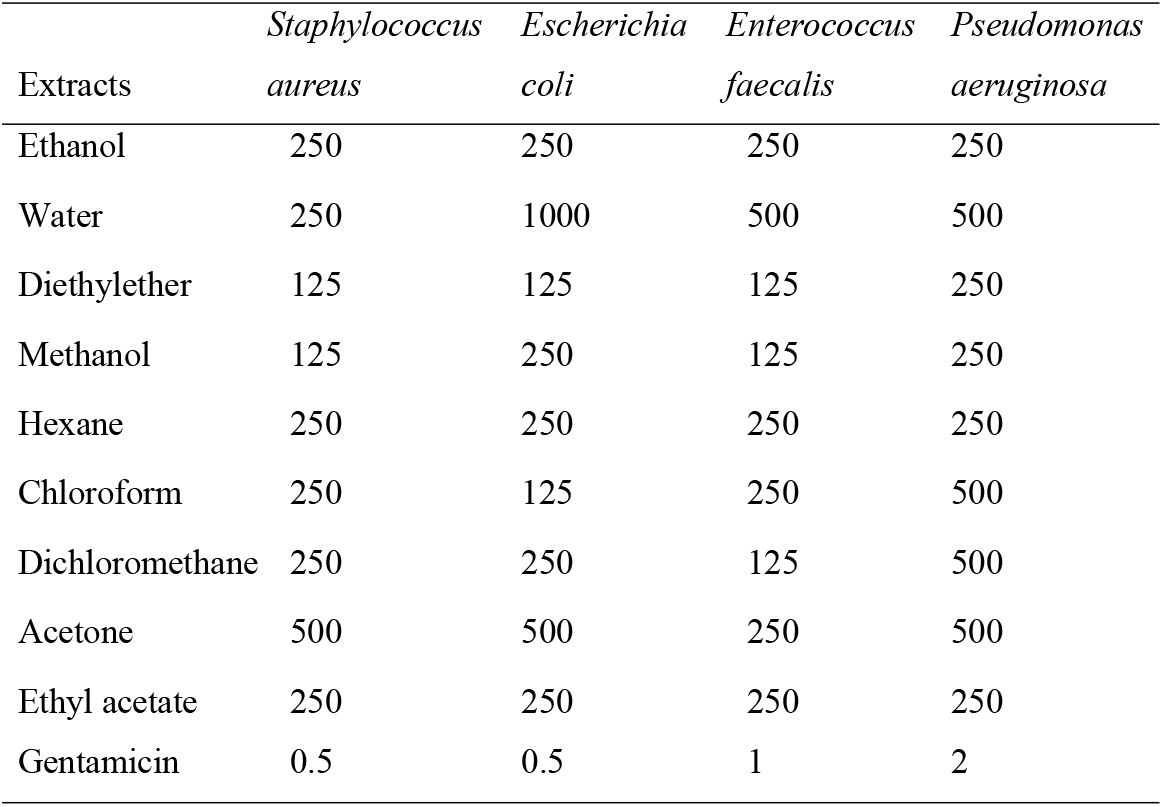
Minimum inhibitory concentration (MIC in µg /mL) values of *Lavandula × intermedia* extract.

| Extracts | *Staphylococcus aureus* | *Escherichia coli* | *Enterococcus faecalis* | *Pseudomonas aeruginosa* |
| --- | --- | --- | --- | --- |
| Ethanol | 250 | 250 | 250 | 250 |
| Water | 250 | 1000 | 500 | 500 |
| Diethylether | 125 | 125 | 125 | 250 |
| Methanol | 125 | 250 | 125 | 250 |
| Hexane | 250 | 250 | 250 | 250 |
| Chloroform | 250 | 125 | 250 | 500 |
| Dichloromethane | 250 | 250 | 125 | 500 |
| Acetone | 500 | 500 | 250 | 500 |
| Ethyl acetate | 250 | 250 | 250 | 250 |
| Gentamicin | 0.5 | 0.5 | 1 | 2 |

**Figure. 1.**
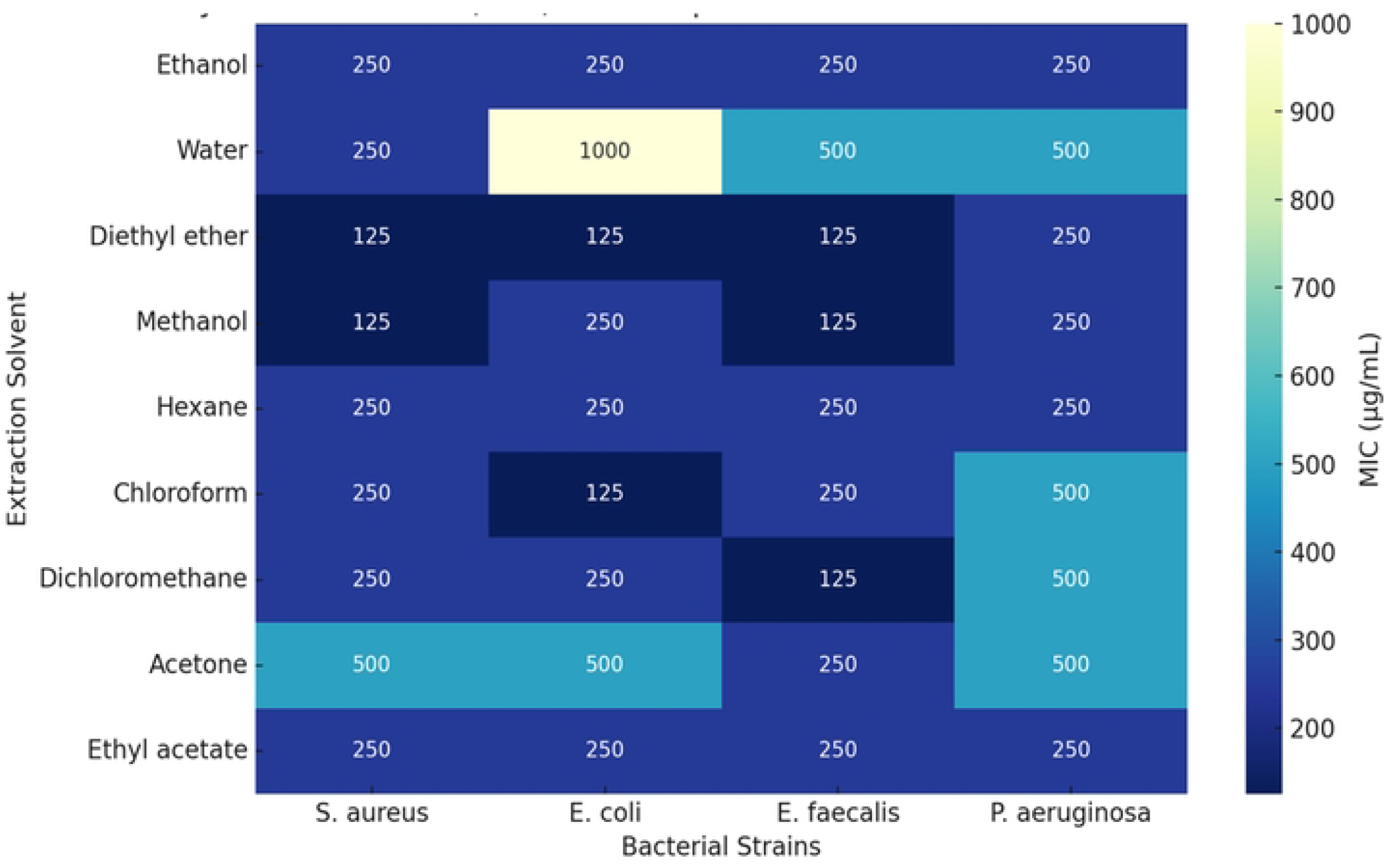

**Figure. 2.**
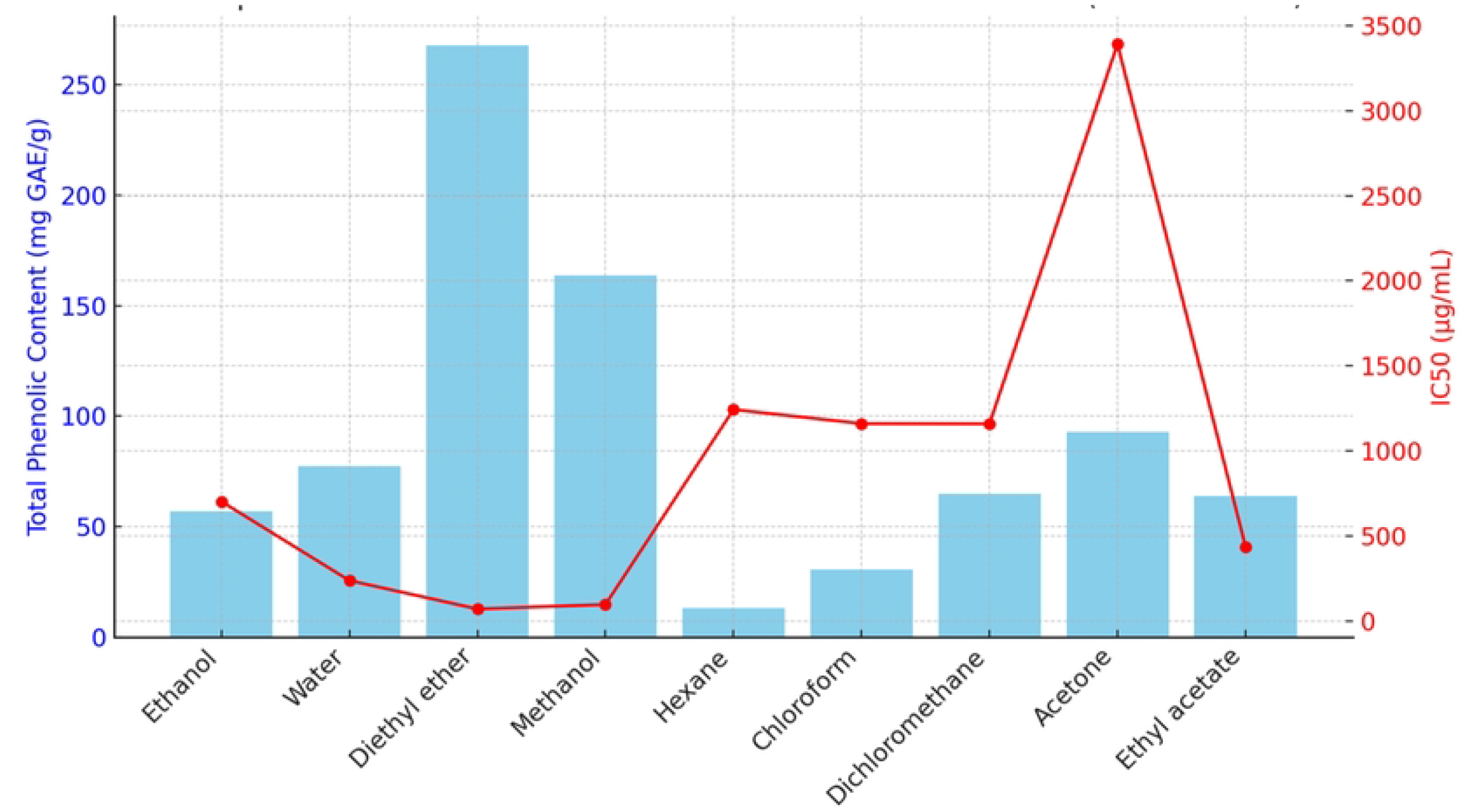

**Figure 3.**
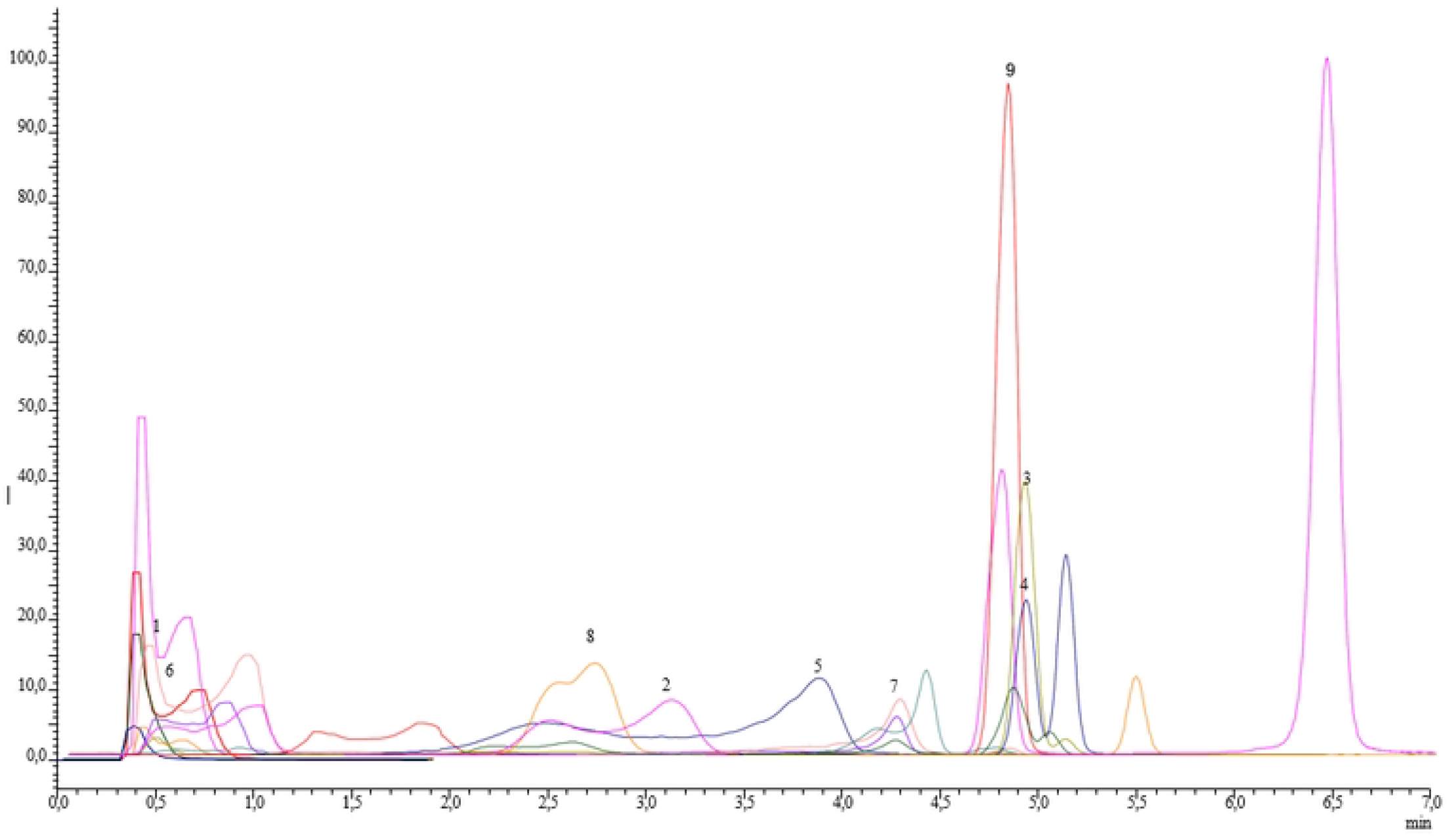
LC-MS/MS chromatograms, 1Fumaric Acid, 2Hydroxycinnamic Acid, 3 Luteolin, 4 Kaempferol, 5 Resveratrol, 6 Protocatochuik Acid, 7 Phloridzin dihydrate, 8 Sabicybic acid, 9 Naringenin.

### Antioxidant activity and total phenolic content

Our findings validate the hypothesis that moderate-polarity solvents maximize bioactive yield. Diethyl ether and methanol extracts significantly outperformed others, yielding the highest TPC (267.65 and 163.8 mg GAE/g, respectively) and the most potent antioxidant activity (lowest IC_50_ values: 72.26 and 98.16 µg/mL) (Table 2). Conversely, solvents at the extreme ends of the polarity scale, such as hexane and water, resulted in minimal TPC and antioxidant capacity. A Pearson correlation analysis revealed a consistent and statistically significant positive correlation between IC_50_ values and MIC values (*p* < 0.05). The strength of the correlation was very strong against Gram-positive bacteria (*r* = 0.887 for *S. aureus* and *r* = 0.771 for *E. faecalis*). This observation is consistent with previous reports by Danh et al. (2013) and Lahkimi et al. (2020), which showed a direct correlation between phenolic content and antioxidant potential in *Lavandula* extracts.

**Table 2.** Antioxidant Capacity and Total Phenolic Content Values of *Lavandula × intermedia*.

| Extracts | DPPH IC<sub>50</sub><br>μg/mL | Total Phenolic Content<br>μg GAE/g Dry extract |
| --- | --- | --- |
| Ethanol | 703.4 | 56.95 |
| Water | 238.93 | 77.35 |
| Diethylether | 72.26 | 267.65 |
| Methanol | 98.16 | 163.8 |
| Hexane | 1,243.66 | 13.3 |
| Chloroform | 1,161.33 | 30.75 |
| Dichloromethane | 1,160.33 | 64.75 |
| Acetone | 3,390 | 92.9 |
| Ethyl acetate | 438.75 | 63.75 |

### Phenolic compound profile of *Lavandula* extract

The phenolic composition was determined by LC–MS/MS analysis (Table 3). A total of nine phenolic compounds were identified and quantified. Among them, fumaric acid was the most abundant compound (1.33 ± 0.08 µg/g), followed by resveratrol (0.67 ± 0.02 µg/g) and hydroxycinnamic acid (0.57 ± 0.02 µg/g). The LC–MS/MS analysis identified these predominant constituents, which are widely reported in the literature to possess both radical scavenging capabilities and direct antimicrobial mechanisms (Hwang and Lim 2015; Tayeh 2025). Notably, this study fills a gap in the literature, as no prior research has evaluated *Lavandula × intermedia* using these specific extract types and the microdilution method.

**Table 3.** Phenolic compound content of *Lavandula × intermedia* extract determined by LC-MS/MS. ^a^RT: Retention time, ^b^ R^2^: Coefficient of determination, ^c^ LOD (µg/L): Limit of detection, ^d^LOQ (µg/L): Limit of quantification. Note: Samples were analyzed in triplicate (n = 3) to ensure precision and reliability.

|  | Phenolic Compound | RT<sup>a</sup> (min) | R<sup>2b</sup> | m/z (precursor → product) | Ionization mode | Amount (µg/g) | LOD<sup>c</sup> (µg/L) | LOQ<sup>d</sup> (µg/L) | Major Biological Activity | Reported in Similar Studies |
| --- | --- | --- | --- | --- | --- | --- | --- | --- | --- | --- |
| 1 | Fumaric acid (–) | 0.47 | 0.9980 | 115.00 → 70.90 | Negative | 1.33±0.08 | 2.76 | 9.20 | Antioxidant, metabolic regulator | Detected in *L. stoechas* and *L. angustifolia* methanolic extracts (Zuzarte et al., 2009; Ez zoubi et al., 2020) |
| 2 | Resveratrol (+) | 3.82 | 0.9999 | 229.00 → 135.00 | Positive | 0.67±0.02 | 7.91 | 26.38 | Antibacterial, anti-inflammatory, antioxidant | Previously reported in *L. angustifolia* and *L. × intermedia* (Božović et al., 2017; Koulivand et al., 2013) |
| 3 | Hydroxycinnamic acid (–) | 2.91 | 0.9995 | 163.00 → 119.10 | Negative | 0.57±0.02 | 7.79 | 25.98 | Antioxidant, UV-protective, antimicrobial | Common phenolic acid in *Lavandula* spp. (Basch et al., 2004; Ez zoubi et al., 2020) |
| 4 | Protocatechuic acid (–) | 0.55 | 0.9999 | 153.20 → 108.00 | Negative | 0.14±0.01 | 7.17 | 23.90 | Antioxidant, anti-inflammatory | Found in *L. stoechas* ethanol extracts (Zuzarte et al., 2009) |
| 5 | Kaempferol (+) | 4.94 | 0.9999 | 286.90 → 153.00 | Positive | 0.10±0.01 | 3.92 | 13.06 | Antioxidant, cytoprotective, antibacterial | Reported in *L. angustifolia* and *L. latifolia* (Božović et al., 2017; Ez zoubi et al., 2020) |
| 6 | Luteolin (–) | 4.96 | 0.9999 | 284.80 → 133.00 | Negative | 0.08±0.01 | 6.4 | 21.4 | Antioxidant, neuroprotective | Identified in *Lavandula officinalis* and *L. intermedia* (Koulivand et al., 2013) |
| 7 | Phloridzin dihydrate (–) | 4.21 | 0.9988 | 434.90 → 273.10 | Negative | 0.04±0.01 | 4.34 | 14.45 | Antioxidant, anti-diabetic | Present in hydroalcoholic extracts of *L. stoechas* (Bousta et al., 2020) |
| 8 | Naringenin (–) | 4.87 | 0.9985 | 270.90 → 151.00 | Negative | 0.01±0.01 | 38.5 | 128.2 | Antioxidant, anti-inflammatory | Detected in *Lavandula* spp. methanolic extracts (Ez zoubi et al., 2020) |
| 9 | Salicylic acid (–) | 2.63 | 0.9981 | 137.20 → 93.10 | Negative | 0.03±0.01 | 3.9 | 13.0 | Antibacterial, signaling molecule in stress response | Reported in *Lavandula officinalis* (Basch et al., 2004) |

## Conclusion

This study confirms that the bioactive potential of *Lavandula × intermedia* is heavily dependent on solvent polarity. Moderate-polarity solvents, specifically diethyl ether and methanol, are superior for maximizing both antioxidant capacity and antibacterial efficacy. LC–MS/MS profiling elucidated that this potent activity is driven by a rich composition of phenolic acids, stilbenes, and flavonoids. These findings provide a clear roadmap for industrial and pharmaceutical applications, suggesting that optimized extraction protocols can enhance efficiency and quality control.

## Declarations

### Credit authorship contribution statement Gülcan Gürses

Conceptualization, Investigation, Methodology, Resources, Editing, Writing - Original Draft.

### Data availability statement

The data generated and analyzed during the current study are available from the corresponding author upon reasonable request.

